# HIV-1-induced small T cell syncytia: susceptibility to direct NK cell killing

**DOI:** 10.64898/2026.06.14.732158

**Authors:** Jon P Girard, Emily E. Whitaker, Christen Frandina, Petra Veljkovic, Madeline K. Carr, Menelaos Symeonides, Markus Thali

## Abstract

Intravital imaging studies of HIV-1-infected humanized mice by three independent groups revealed the presence of small T cell syncytia. Although these multinucleated entities represent only a small fraction of infected T lymphocytes, they arise at the earliest stages of infection, are highly motile, and they make frequent contact with potential target cells. Importantly, while such transient contacts allow for virus transfer, they typically do not result in fusion and these exclusively T cell-based syncytia thus do not grow beyond the three- or four-nuclei stage.

Immunofluorescence microscopy analyses, together with initial multiparameter flow cytometric evaluation and proteomic profiling by our lab and collaborators, identified these small T cell syncytia (henceforth referred to merely as syncytia) as a distinct subpopulation of infected cells. Because they appear as soon as any infected cells are detectable and thus likely contribute to early viral spread, they also trigger innate immune responses, including attacks by cytolytic effectors such as natural killer (NK) cells.

Here, we first introduce a newly developed surface split GFP system which allows to unequivocally distinguish syncytia from infected mononucleated cells, and which also enables isolating these entities for in-depth functional analyses. Then, using multiparameter flow cytometry with UMAP clustering and transcriptomic profiling, we document that syncytia represent a subpopulation of infected cells that do indeed clearly differ from both uninfected and infected mononucleated T lymphocytes, particularly also regarding expression of immune-regulatory host factors. Finally, we show that the rate of direct killing by NK cells is substantially higher for syncytia than for infected mononucleated cells, both in 2D suspension co-culture and in a 3D collagen coculture system.

While the fitness costs imposed on syncytia by such increased susceptibility to NK cytolysis during early infection might be compensated for by yet to be determined syncytia functions that (directly) enhance virus spread, we note that irrespective of what such virus spread-enhancing functions may be, the increased vulnerability of syncytia could possibly be exploited for the development of novel antiviral strategies.

## Introduction

Human immunodeficiency virus type 1 (HIV-1) dissemination critically depends on migration of infected T cells. Unexpectedly, a minor fraction of the infected T cells exist as small syncytia, containing up to four nuclei [1–3]. Importantly, these entities are not precursors for larger, macrophage- or dendritic cell-based syncytia which can be observed in late stages of virus dissemination/pathogenesis. Rather, and as shown in three independent intravital imaging studies, they are present already at the earliest stages of infection (for a review, see [4], also [3]. Our analyses in physiologically relevant *in vitro* settings [5] for the first time documented that these small T cell syncytia (henceforth referred to merely as syncytia) can transfer virus to uninfected cells, suggesting, as also commented on in the above cited review [4], that they can directly contribute to virus dissemination. Further, with another previous study in our lab, we started documenting that the surface protein profile of syncytia differs substantially from the profile of infected mononucleated cells [6]. Together with more recent, preliminary investigations that included quantitative migratory behavior analysis (using Migrate 3D, an in-house developed novel software;[7], multiparameter flow cytometric evaluation, immunofluorescence microscopy analyses, and initial proteomic profiling (in collaboration with the Matheson group at U Cambridge), establish that syncytia represent a distinct subpopulation of HIV-1-infected cells.

Given that syncytia arise immediately after infection, these entities, like infected mononucleated cells, need to deal with the host’s innate immune response, which includes attacks by cytotoxic effectors such as natural killer (NK) cells. Recent studies on how NK cells deal with HIV-1-infected cells have focused mostly on antibody-dependent cellular cytotoxicity (ADCC). NK cells, however, which constitute 5 to 15% of peripheral blood mononuclear cells, act against HIV-1-infected cells already very early upon infection, not only through release of cytolytic granules containing perforin [8] [9], but also via secretion of cytokines and other signaling molecules [8, 10–12]. Such early interactions clearly contribute to disease outcome in HIV-1 infected individuals [13]. That NK cells play a pivotal role also in (early) immune control was very strongly suggested with a study that reported on virus-exposed but uninfected individuals [14] and also recently with a study [15] which showed that two HIV-1 controllers had a distinct proliferative, cytotoxic subset of NK cells that appeared very early in acute HIV-1 infection, i.e., before appearance of CD8^+^ T cell responses.

The outcome of interactions between HIV-1 infected cells and NK cells depends on multiple factors, including HIV-1 Nef-induced major histocompatibility complex class 1 (MHC-I) downregulation, which, while providing protection of infected cells against antigen-dependent CTL attacks, exposes them to antigen-independent (i.e. innate) NK cell-mediated direct killing [16–18]. In addition to recognition of MHC-I on target cells, expression of other inhibitory receptors on NK cells together with activating ligands produced by infected cells, determines whether NK cells will secrete inflammatory cytokines and/or also directly kill infected cells [12]. Consequently, with our current investigations of the properties of syncytia, we focused on factors involved in regulating response by NK cells as well as on actually measuring that response. Given that the presence of extracellular matrix is known to affect many cellular properties, including NK cell cytotoxicity [19], and also as unpublished data from our lab suggest that syncytia are able to utilize 3D environments to move and make cell-cell contacts differently than infected mononucleated cells, we set out to measure syncytia-NK cell interactions both in a more traditional suspension culture system as well as in a 3D collagen coculture system.

## Results

### Surface split GFP system for syncytium identification and isolation

HIV-1-induced, small T cell syncytia most likely are the result of an occasional failure to resolve the virological synapse upon transfer of viral particles, i.e., when infected and target T cells, rather than separating, fuse with each other, as depicted in Fig 1A. To start characterizing these unique and relatively rare cells, we developed a novel surface split GFP system which allows one to unequivocally distinguish syncytia from infected mononucleated cells, and which also enables physically isolating these entities for highly specific and functional analyses, including for transcriptomics.

**Figure 1.**
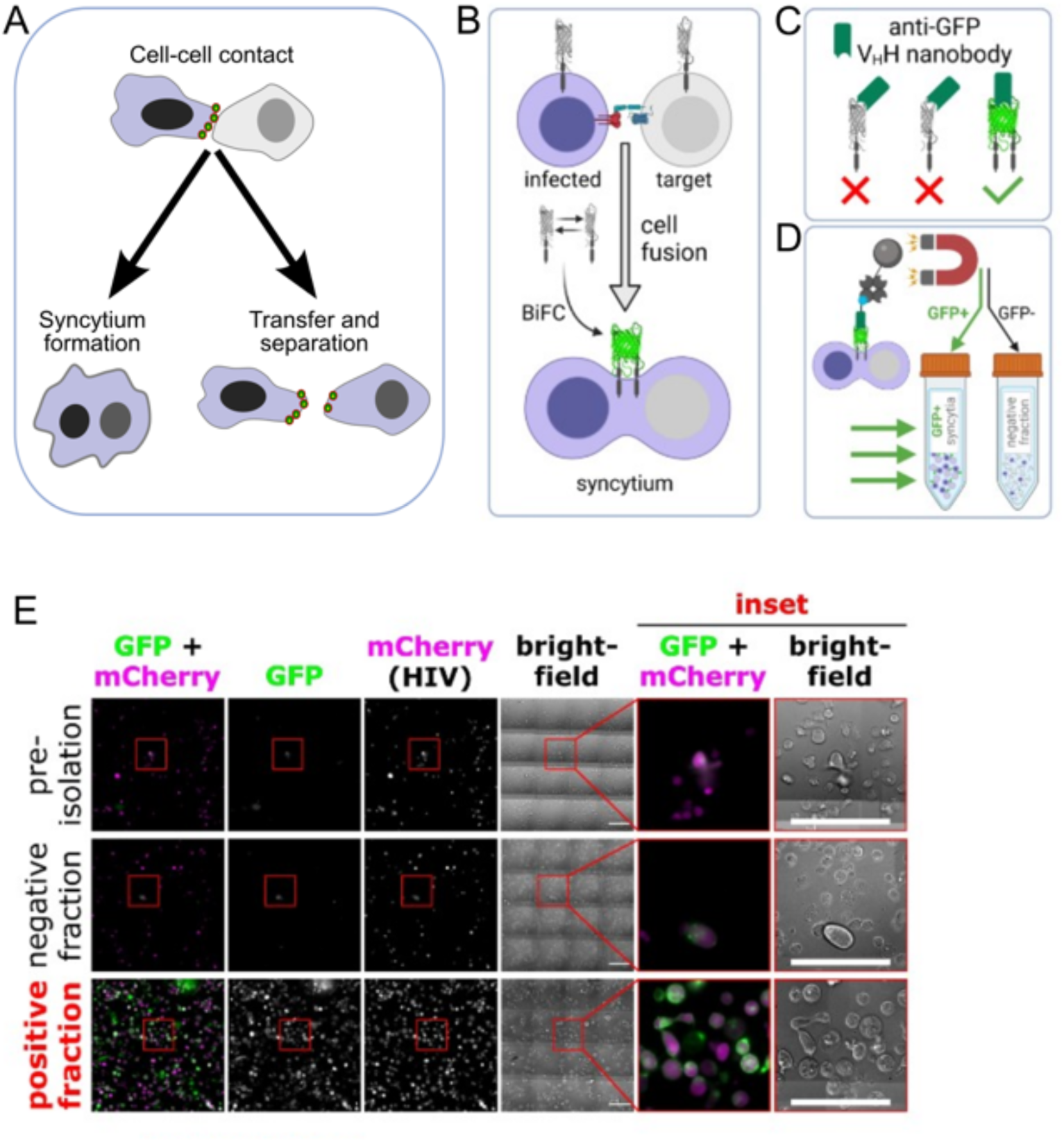
Syncytia isolation procedure. **(A)** Syncytia formation at the virological synapse during HIV transmission events **(B)** GFP2-Display-expressing cells infected with HIV-1 NL-CI (mCherry reporter NL4-3, from B.K. Chen) are co-cultured with target cells stably expressing GFP1-Display. Upon HIV-1-induced fusion between these two cell populations, syncytia form and, upon bimolecular complementation between GFP1 and GFP2, bear assembled whole GFP on their surface. **(C)** Surface whole GFP is then detected using a highly specific biotinylated anti-GFP nanobody (Chromotek #gtb-250) that does not bind the individual split GFP components. **(D)** After labeling the co-cultures with the biotinylated anti-GFP nanobody, which will only bind the syncytia in the culture, syncytia are positively isolated using an anti-biotin magnetic cell isolation kit (StemCell). This strategy can be used in any cell line and with any virus strain. **(E)** Microscopic evaluation of syncytia isolation procedure performed using NL-CI-infected A3.01 T cells and A3R5 target cells. Scale bars = 100 μm.

Fig 1B shows how fusion of cells that express either portion of split GFP (GFP1 or GFP2, respectively) results in bimolecular complementation and thus the expression of whole GFP on the surface of these newly formed syncytia and can then serve as marker for these entities.

Figs 1C-E illustrate how the same bimolecular complementation system can be used to isolate syncytia. At the beginning of this newly developed isolation strategy, A3.01 and A3R5.7 cells are separately transduced with constructs that allow for surface expression of either portion of the split GFP, thus forming A3.01-G2D and A3R5-G1D cells, respectively, which upon cell-cell fusion will assemble full GFP. To produce syncytia, A3.01-G2D cells are first spinoculated with VSV-G-pseudotyped NL-CI^JRFL Env^ (an mCherry reporter strain of HIV-1 derived from NL4-3 and bearing an R5-tropic Env). Two days later, HIV-1-infected cells are isolated using negative selection for CD4 and HLA-A which are both downmodulated upon HIV-1 infection. Isolated infected cells are then cocultured with A3R5-G1D cells in the presence of reverse transcriptase inhibitor to prevent viral transmission, thus allowing all uninfected cells to serve as potential fusion targets rather than viral transmission targets.

Following 40 h of coculture, syncytia can be immunomagnetically isolated using a highly specific anti-GFP nanobody (that recognizes only fully assembled GFP but cannot bind to either split portion of it) (Fig 1C-D; quantification is shown in Fig 1E). Besides allowing us to isolate syncytia, whole cocultures can also be used for various downstream experiments, where infected mononucleated cells will be mCherry+/GFP-, syncytia will be mCherry+/GFP+, and uninfected cells will be mCherry-/GFP-. The purity of the resulting isolated syncytia is thus easily evaluated by flow cytometry, and in our hands has consistently been near-100% (data not shown). To our knowledge, this is the first report of a syncytia isolation system (not only for HIV-1-induced syncytia, but in any context).

### Syncytia represent a distinct subset of HIV-1-infected cells

2While already our initial *in vitro* analyses of these syncytia, combined with additional evaluation of intravital imaging data generated in one of the three humanized mouse studies [5], strongly suggested that these entities differ substantially from infected mononucleated T cells, more recent quantitative analyses of syncytia migration and motility provided further evidence for them being unique entities (data not shown). While some of the observed changes may be due to altered ratios of host and viral factors, recent studies have documented that cell-cell fusion per se, i.e., independent of viral infections, can cause major transcriptional reprogramming [20], equivalent to a developmental switch. We thus sought to further profile our HIV-1-induced syncytia, focusing on expression of surface host factors (Figs 2A and B) and the transcriptome (Fig 2C), and comparing these entities to uninfected T lymphocytes and to infected mononucleated cells. Given their presence at the earliest stages of HIV-1 infection in humanized mice [1–3] and the likelihood that syncytia will interact with early innate immune effector cells, including NK cells, we thus focused part of these analyses on the presence (or absence) of immune regulatory factors.

**Figure 2.**
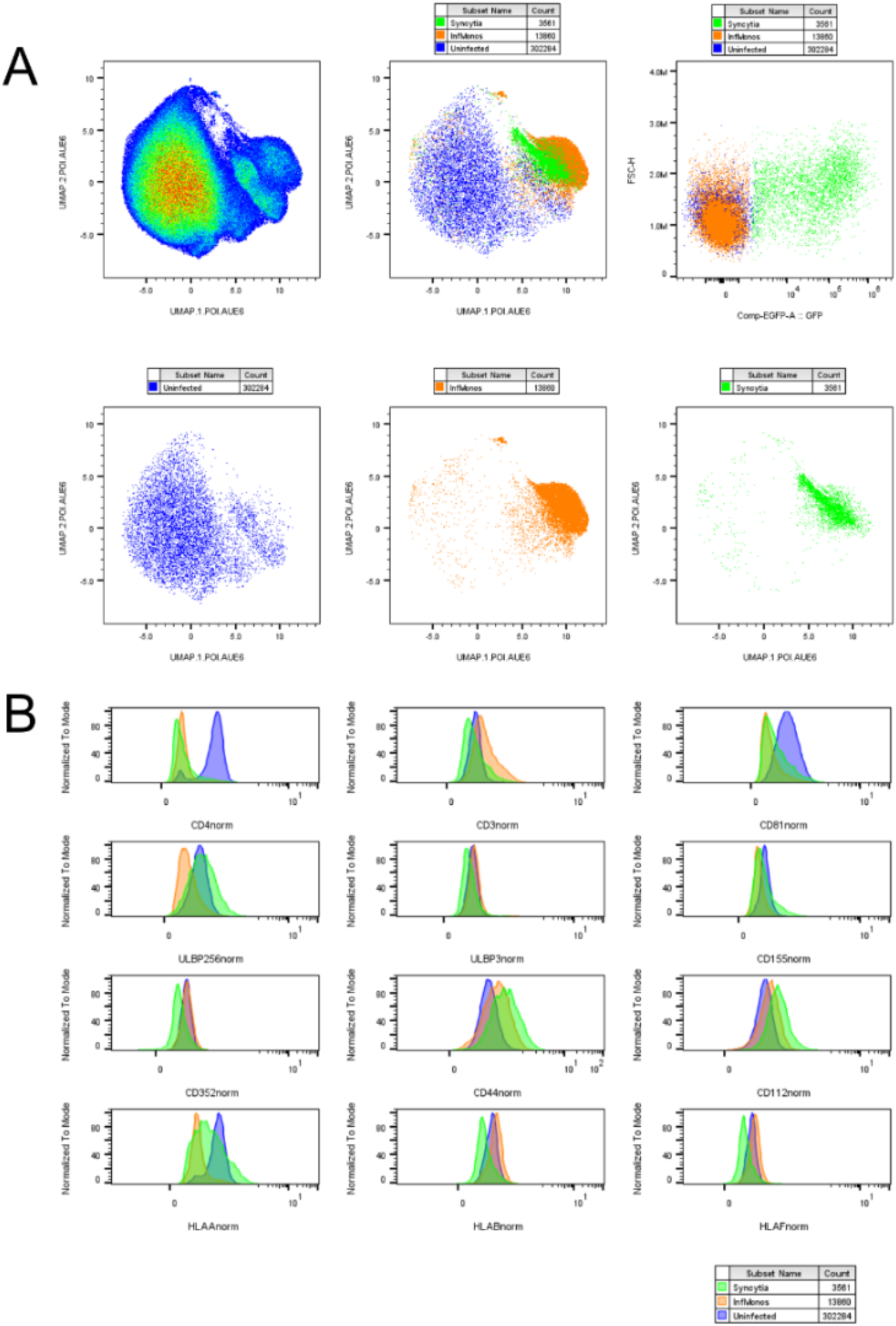
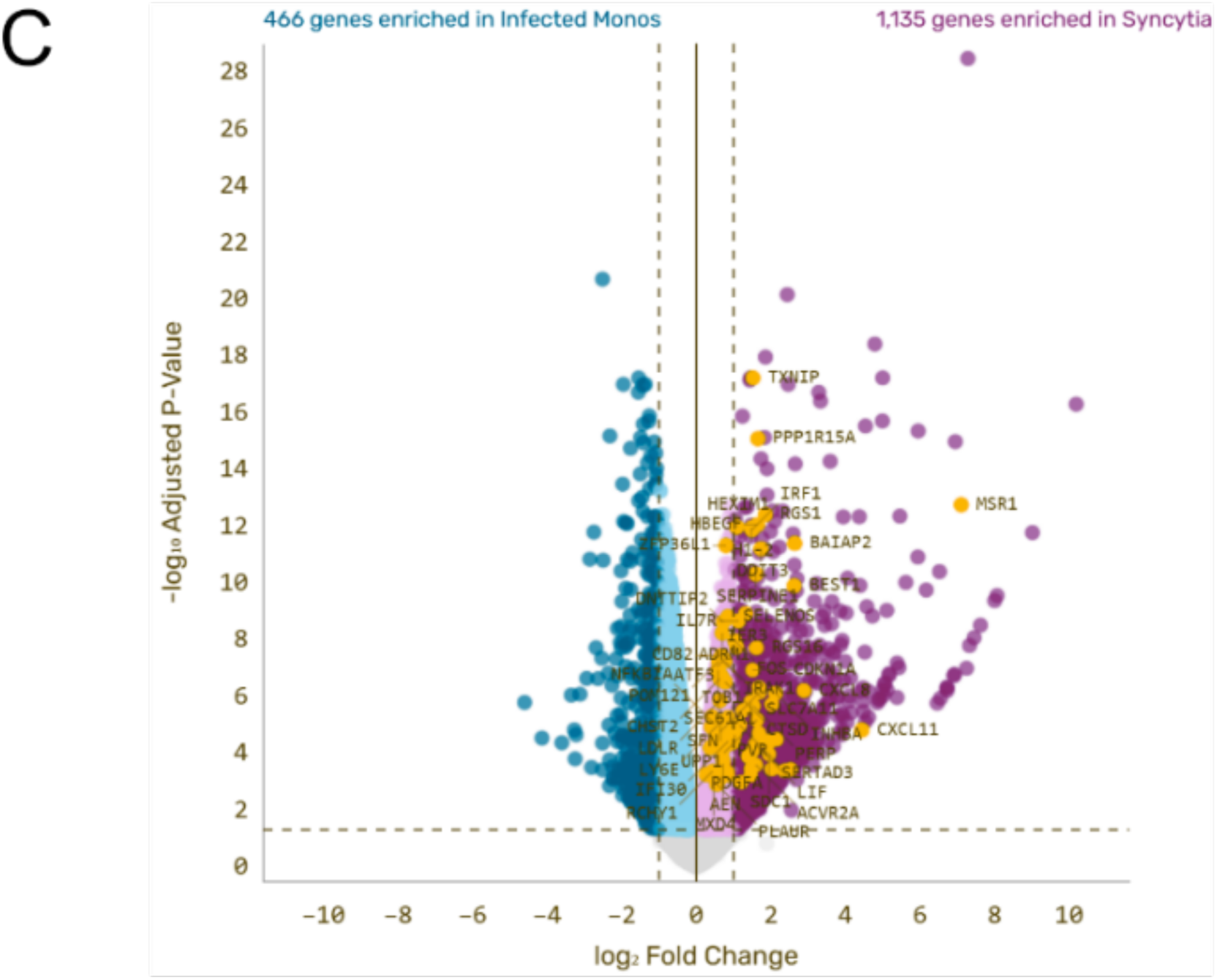
Flow cytometric and RNAseq evaluation of syncytia. Co-cultures of GFP2-Display-expressing HIV-1-infected (NLCI JRFL Env) A3.01 cells and GFP1-Display- and CCR5-expressing A3R5.7 target cells were surface-stained for the proteins shown and analyzed by spectral flow cytometry on a Cytek Aurora instrument. Live cells were identified and gated for infection using the mCherry reporter and syncytia were identified as mCherry+GFP+. The values corresponding to each surface protein of interest were normalized to cell size by dividing by the forward scatter height (FSC-H) parameter. **(A)** UMAP analysis was performed on the resulting datasets, excluding mCherry and GFP so as to not bias the clustering of cell subsets. In other words, the UMAP algorithm was given no information on which cells were uninfected, infected mononucleated, or syncytia. **(B)** Histograms of each surface protein stained for in the panels, with each cell subset shown as separate overlays. (**C**) Differential gene expression analysis of isolated syncytia VS and isolated infected mononucleated cells. Shown in yellow are genes involved in inflammatory responses and apoptosis. (RNAseq analyses performed by Plasmidsaurus for **C**).

Using multiparameter flow cytometry with UMAP clustering, we first show that syncytia make up their own unique population of infected cells (Fig 2A). Host factors that contribute to the distinct clustering of syncytia include PVR/CD155 and NTBA/CD352 which are NK cell activating ligands that promote direct killing by NK cells (Fig 2B) [12]. With the goal of looking beyond differential expression of host factors at the surface of the different populations of infected cells, syncytia and infected mononucleated cells (isolated as described in Fig 1 and subsequently lysed) were then subjected to bulk RNAseq analysis. As shown in Fig 2C, genes that are differentially expressed in syncytia include many that are involved in inflammatory signaling and apoptosis (highlighted in yellow). Note, that PVR (CD155), whose surface (protein) expression is documented in Fig 2B, has also higher RNA expression in syncytia.

Taken together, flow cytometry and RNAseq data shown in Fig 2 lead us to hypothesize that syncytia are more susceptible to NK cell direct killing than infected mononucleated cells. Given the presence of syncytia early in HIV-1 infection, and also given that early NK cell activities are an immediate response to HIV-1 exposure [14], we decided to investigate if syncytia are relatively more susceptible to direct killing by NK cells.

### Syncytia are more susceptible to NK cell-mediated killing than infected mononucleated cells

#### a) Susceptibility in 2D suspension coculture

NK cell direct killing is an innate immune response that is controlled by a diverse mixture of activating and inhibiting receptors [12, 18]. As shown above, multiple activating ligands such as PVR/CD155 (protein and RNA) Nectin-2/CD112, and ULBP2/5/6 all have an increased presence on syncytia (Fig 2B). PVR and Nectin-2 are ligands for the activating NK cell receptor DNAM-1 and ULBP2/5/6 are ligands for the activating NK cell receptor NKG2D.

To study the susceptibility of syncytia to NK cell direct killing we utilized our cell-cell fusion reporter described in Fig 1 and the NK cell line KHYG-1. These cells express the general direct killing receptors and are easily cultured in the lab. The experimental setup for these killing experiments is depicted in Fig 3A. Briefly, syncytia-producing cocultures (as described earlier) were set up and then mixed with Celltrace Far Red-dyed KHYG-1 cells for 6 h. In addition to the cocultures, we also set up non-coculture controls consisting of target cells and KHYG-1 cells cultured separately for the same period of time and combined right before staining. Cells were then harvested and stained for viability using Live/Dead Blue, fixed and used for flow cytometry. Fig 3B depicts the gating scheme used, which allows us to identify all populations of cells within our cocultures. Percentage of killed cells is determined by normalizing the number of each type of target cell to the number of KHYG-1 cells in that sample, and dividing these normalized counts by the analogous normalized counts in the non-cocultured controls. Our results show that syncytia are more susceptible to NK cell direct killing in these 2D cell culture conditions.

**Figure 3.**
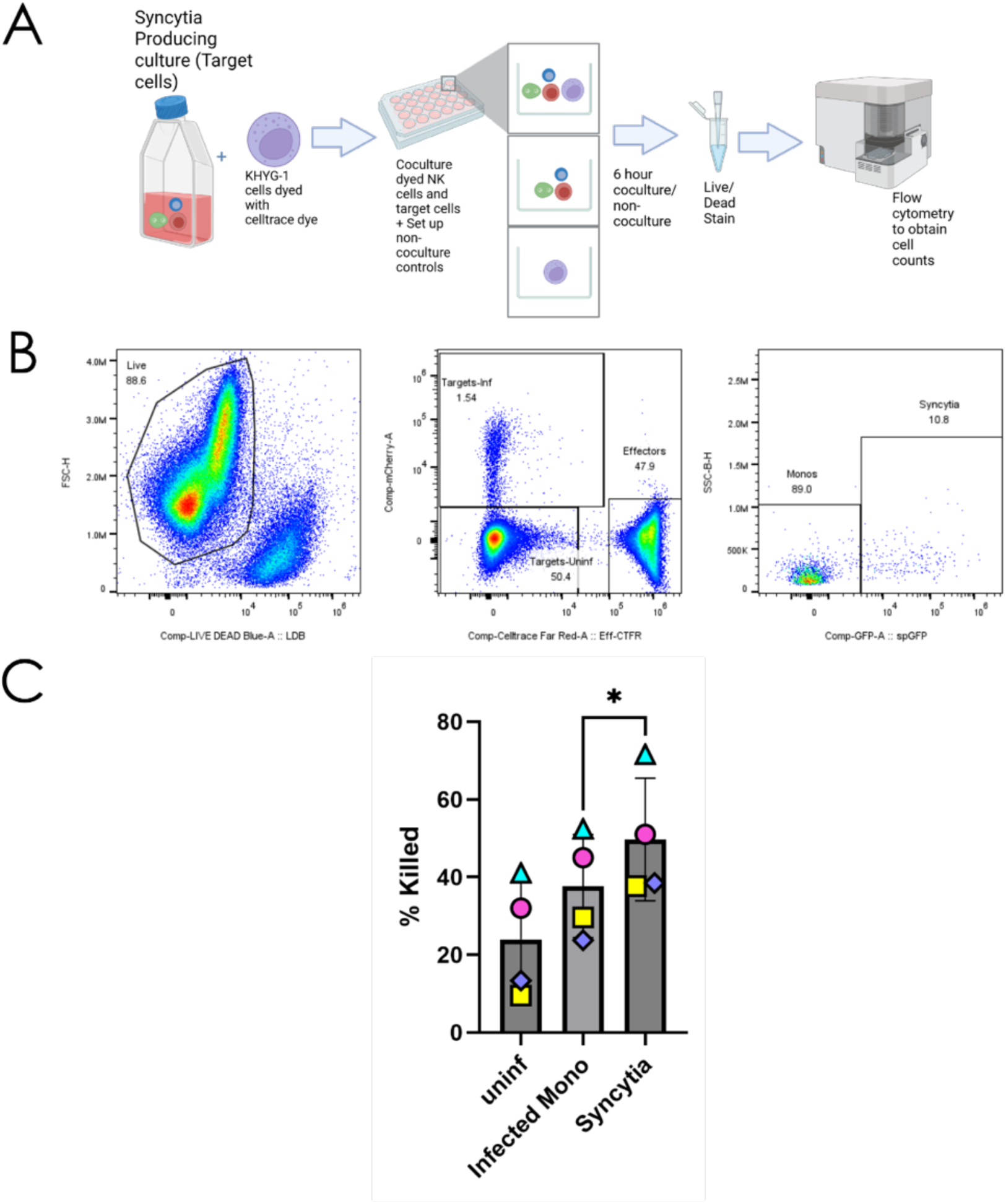
NK cell killing in 2D. (**A**) Target cells (syncytia, infected mononucleated cells and uninfected T cells) were cocultured with dyed KHYG-1 cells (NK cell line) for 6 hrs in 2D cell culture. In addition, non-cocultured controls (target cells or KHYG-1 cells cultured alone) were also plated. After coculture, cells were harvested directly. Cells were stained for viability using Live/Dead blue, fixed and then used for flow cytometry to measure killing. (**B)** Gating scheme for killing experiments. (**C**) 2D coculture killing data. % killing was determined by normalizing each target cell count to the effector cell count and dividing the cocultured conditions by the non-cocultured controls. * = P<0.05

#### b) Susceptibility in a 3D collagen coculture system

Given that our (so far unpublished), semi-quantitative analyses of the motility/migratory capabilities of syncytia suggest that their interactions with potential target cells, when analyzed in 3D cell culture systems, i.e. in the presence of extracellular matrix, differ from those of infected mononucleated cells, we went on to measure killing of syncytia in such a system. To do that, we used the same general setup as for the 2D version of the experiment except that the cells were embedded in 3D collagen gels (Fig 4A) which were then dissolved using collagenase to release the cells after coculture. Following the same gating scheme, we quantified cell killing and found that, like in the 2D coculture system (Fig 3C), syncytia are more prone to be killed by the NKs than infected mononucleated cells. Interestingly, and to some extent against our expectations, syncytia were also found to be considerably more susceptible to killing in 3D environments than they are in 2D cell culture system.

**Figure 4.**
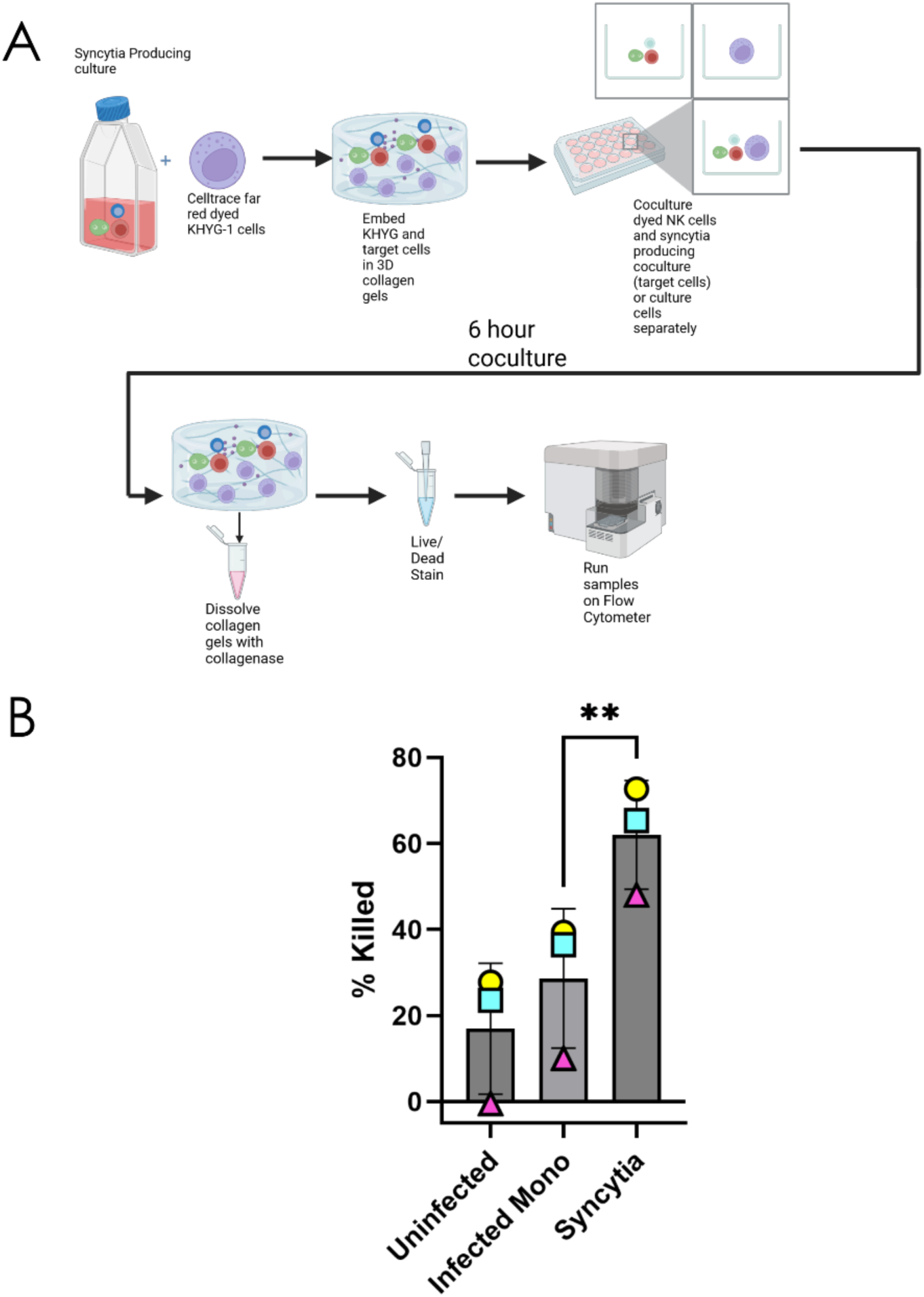
NK cell killing in 3D. (**A)** Target cells (syncytia, infected mononucleated cells and uninfected T cells) were cocultured with dyed KHYG-1 cells (NK cell line) for 6 hrs in 3D collagen gels. In addition, non-cocultured controls (target cells or KHYG-1 cells cultured alone) were also plated in 3D collagen gels. After coculture, cells were harvested after collagen gels were dissolved using collagenase. Cells were stained for viability using Live/Dead blue, fixed and then used for flow cytometry to measure killing. (**B)** 3D collagen gel killing data. % killing was determined by normalizing each target cell count to the effector cell count and dividing the cocultured conditions by the non-cocultured controls. ** = P<0.01.

## Discussion

While we have previously reported on HIV-1-induced syncytia displaying distinct motility [5] and altered surface expression (relative to infected mononucleated cells) of proteins that prevent cell-cell fusion (and thus presumably continued growth of syncytia) [6], here we now present initial results of our investigations of differential expression of immune-regulatory proteins in syncytia (compared to infected mononucleated cells) and of how that affects their interactions with NK cells. Most importantly, data presented here (in Fig 2) confirm what we have already seen in (still preliminary; data not shown) analyses of their proteomes: syncytia and infected mononucleated cells are clearly distinct entities! While they share some of the up- or down-regulation of host factors that are regulated by HIV-1 accessory proteins, the majority of changes seen in the RNAseq analysis apparently are not caused by the infection per se but likely are the result of cell-cell fusion. Also, while some host factors that are downregulated in infected mononucleated cells are further downregulated in syncytia, surface expression of others, including e.g. of CD4, appear to be (relatively) rescued in syncytia. Thus, HIV-1 Env-induced T cell-T cell fusion, comparable to what was reported for other cell types [20], reprograms these lymphocytes, changes their identity, and equips them with novel properties. While some of them may be beneficial (for the infected cell), others may weaken them, at least as far as one can measure within currently available experimental systems: as shown here (Figs 3 and 4), syncytia are more readily killed by NK cells than infected mononucleated cells. While it remains to be investigated how the reprogramming renders syncytia more prone to be killed by NK cells (presumably via cytotoxic granules), our RNAseq analysis and surface profiling by flow cytometry suggests that relatively augmented expression of distinct activating ligands, including PVR, Nectin-2, and ULBP2/5/6 (Fig 2B & C), may lead to increased cytotoxic NK action. Additionally, since syncytia have influx of surface host proteins from uninfected cells, this may reverse some of the cell surface remodeling induced by HIV-1 accessory proteins Nef and Vpu in infected mononucleated cells, thus rendering them less prone to be attacked by NK cells.

Determining exactly what renders syncytia more prone to be killed by NK cells will be interesting also as it may help guiding investigations aimed at explaining why killing is enhanced under 3D conditions. As mentioned, this result was somewhat unexpected as our preliminary quantitative analysis of motility and migration capabilities of syncytia suggested to us that these entities might be better at evading approaching immune cells, particularly when they can move on extracellular matrix “tracks”, i.e., in a 3D environment. Future studies aiming at providing mechanistic explanation of the increased killing will need to include NK cells, as the presence of those cells may alter syncytia motility, e.g. through delivery of specific inhibitory signals. Alternatively, infected mononucleated cells may be more able to withstand cytolytic attack, comparable to how HIV-1-infected macrophages have recently been shown to withstand NK attacks and direct killing relatively better than CD4^+^ T cells as they can skew NK responses toward production of TNF-α over degranulation by NK cells [9].

Irrespective of the potential mechanistic explanation, we can think of (at least) two reasons for why HIV-1 “tolerates” such increased vulnerability of syncytia: 1) Given that, as suggested with Fig 1A, virus transmission at cell-cell contact sites is mostly followed by separation of producer and target cells, the occasional fusion events could simply be an unwanted byproduct of an otherwise very efficient transmission process, a byproduct that can be tolerated as the vast majority of infected cells are/remain mononucleated (due to established barriers to cell-cell fusion at the virological synapse, discussed in [21] and thus are less prone to be attacked and killed by NK cells; 2) The formation of a small number of syncytia might even be desirable as these entities might serve as decoys and/or might be sacrificing themselves (in Arnold von Winkelried fashion) so that infected mononucleated cells can spread the virus more efficiently, less hindered by cytolytic immune cells.

We should like to suggest, finally, that the increased vulnerability of syncytia could possibly be exploited for the development of novel anti-viral strategies: by enhancing cell-cell fusion at the virological synapse and thus augmenting the number of newly formed syncytia, one could generate “easy targets” for elimination by the immune system. Enhancing of HIV-1 Env-induced fusion could be achieved through pharmacological dysregulating of Aurora Kinase B [22] or through inhibition of phospholipid scramblase 1 (PLSCR1) [23]. Such enhancement of NK cells’ “natural” antiviral activities would be comparable to how CD4 mimetic molecules (which open up distinct antibody binding sites in HIV-1 Env) are being evaluated as tools to enhance antibody dependent cellular cytotoxicity (ADCC) [24]. Indeed, given that CD4 surface expression is relatively restored in syncytia (as discussed above, see Fig 2B) and may interact there with Env, it would seem reasonable to test if syncytia are “naturally” sensitized to ADCC.

## Author Contributions

J.P.G., M.S., and M.T. conceived and designed the experiments. J.P.G. and M.S performed the experiments and analyzed the results, with contributions by E.E.W., P.V., C.F., and M.C. for designing/optimizing various parts of the project including the split GFP fusion reporter, syncytia production, isolation and 3D collagen gels. J.P.G., M.S. and M.T prepared the figures. J.P.G., M.S., and M.T. wrote and edited the manuscript.

## Acknowledgments

The flow cytometry data we presented were obtained at the Harry Hood Bassett Flow Cytometry and Cell Sorting Facility, Larner College of Medicine, University of Vermont. David Evans generously gifted the KHYG-1 cell line. Ben Chen provided the mCherry reporter HIV-1 virus.

## Materials and Methods

### Cell lines and Cell Culture

HEK293T cells were maintained in DMEM (Corning #10090CV) supplemented with 10% fetal bovine serum (FBS; Cytiva #SH3091003) and 1% penicillin-streptomycin (P/S; Gibco #15140122). KHYG-1 cells were grown in RPMI-1640 (Cytiva #SH30027FS) supplemented with 10% FBS, 1% P/S, and 100 U/ml IL-2 (StemCell Technologies #78036.2). A3.01 cells (NIH HIV Reagent Program #ARP-166) were maintained in RPMI-1640 supplemented with 10% FBS and 1% P/S. A3R5.7 cells (NIH HIV Reagent Program #ARP-12386) were maintained in RPMI-1640 supplemented with 10% FBS and 1 mg/mL G418 (InvivoGen #ant**-**gn-2). All cells were maintained at 37°C in 5% CO_2_ atmosphere with 95% humidity.

A3.01-G2D cells were generated by transduction of A3.01 cells with a custom-generated lentiviral vector (pCVL-UCOE-SFFV, Blasticidin resistance marker) for expression of the C-terminal fragment of split-GFP (GFP2) inserted into the ectodomain of the pDisplay construct (Invitrogen #V66020). A3R5-G1D cells were generated by transduction of A3R5.7 cells with two custom-generated lentiviral vectors (pCVL-UCOE-SFFV, Blasticidin resistance marker; and pLX301, Puromycin resistance marker) both expressing the N-terminal fragment of split-GFP (GFP1) inserted into the ectodomain of the pDisplay construct. A3.01-G2D cells were grown in RPMI-1640 supplemented with 10% FBS and 4 μg/mL Blasticidin S (Gold Biotechnology Inc #B800100). A3R5-G1D cells were grown in RPMI-1640 supplemented with 10% FBS, 4 μg/mL Blasticidin S, 1 mg/mL G418 (Invivogen #ant**-**gn-2) and 500 ng/mL Puromycin (Gibco #A11138-03). Cells were routinely checked for mycoplasma by PCR; no positivity was detected at any time.

### Virus production

Viruses were produced in HEK293T cells by co-transfection with NL-CI^JRFL Env^ and pVSV-G using calcium phosphate precipitation. Virus-containing supernatants collected 96 h post-transfection were cleared of cell debris by centrifugation at 2000 rcf for 10 min, filtered through a 0.45 μm filter, mixed at a 3:1 ratio with 4X buffered polyethylene glycol (PEG) solution (40% PEG-8000, 1.2 M NaCl, pH 7.0-7.2 in ddH_2_O), and rotated for 4 h at 4°C. After rotation, PEG-virion complexes were pelleted at 1600 rcf for 1 h at 4°C, the supernatant was discarded, and pellets were resuspended in complete RPMI medium at 1/25^th^ of the initial volume of supernatant. These 25X concentrates were aliquoted and stored at -80°C.

### HIV Infection

A3.01-G2D cells were infected with VSV-G-pseudotyped NL-CI^JRFL Env^ by spinoculation as follows: Cells were pelleted and resuspended at 1x10^7^ cells/mL in complete RPMI medium containing 5 μg/mL polybrene (hexadimethrine bromide; Sigma 107689-10G) and 25X-concentrated virus stock (at a dilution empirically determined to infect ∼15% of the cells). After a 15 min incubation at 37°C/5% CO_2_, cells were centrifuged at 1200 rcf for 90 min at 37°C, then transferred into fresh complete RPMI medium at a density of 5x10^5^ cells/mL. Because an R5 Env-containing virus was used in CCR5-negative cells, infected cells could not propagate the infection among them or form syncytia.

### HIV-1-infected cell enrichment and syncytia production

HIV-1-infected cells were enriched by immunomagnetic depletion of CD4- and HLA-A-expressing cells. 48 h post infection, infected cells were counted, pelleted, and re-suspended in isolation buffer (IB; PBS + 2% FBS + 1 mM EDTA) at 5x10^7^ cells/mL containing 1.25 μg/mL anti-human CD4 antibody (clone OKT4; Biolegend #317402) and 1 μg/mL anti-human HLA-A antibody (clone 108-2C5; Novus Biologicals #NBP2-45320-0.1mg). Cells were rotated at 4°C for 30 min, washed with 10 volumes of ice cold IB, and resuspended in ice cold IB at 1x10^7^ cells/mL. DynaBeads Goat Anti-Mouse IgG (Invitrogen #11033) were added at 85 μL per 1x10^7^ cells, and cells were rotated at 4°C for 30 min. The volume was brought up to 10 mL with ice cold IB, and cells were placed in a magnetic field for 5 minutes. Buffer containing unbound (CD4- and HLA-A-negative) cells was decanted into a collection tube, collected cells were pelleted, counted, pelleted again, and resuspended at 1x10^6^ cells/mL in complete RPMI medium containing 2 μM efavirenz (HIV Reagent Program #HRP-4624).

Enriched HIV-1-infected cells were cocultured with uninfected A3R5-G1D cells at a 1:1 ratio for 40 h. This was done in the presence of 2 μM efavirenz to prevent transmission while allowing formation of syncytia, which was now possible with the use of R5 Env because A3R5 cells express CCR5. Upon cell-cell fusion, G1D and G2D assemble into whole GFP on the surface of syncytia.

### Syncytia isolation

Syncytia-producing cocultures were counted, pelleted, and resuspended in cold IB containing 20 μg/mL biotinylated anti-GFP nanobody (Chromotek #gtb-250) to label syncytia. This nanobody was extensively validated to only bind whole assembled GFP and not either of the split-GFP components alone (data not shown). Cells were then rotated at 4°C for 30 min, pelleted, and resuspended in IB at 5x10^7^ cells/mL. Labeled syncytia were magnetically isolated and released from isolation beads using the EasySep Release Human Biotin Positive Selection Kit (StemCell #17653). Isolated cells were washed three times after isolation to remove any remaining magnetic beads.

### RNAseq preparation and analysis

1x10^5^ isolated syncytia, isolated infected mononucleated A3.01-G2D cells, uninfected A3.01-G2D cells, and uninfected A3R5.7-G1D cells (all in duplicate) were lysed for 30 min using DNA/RNA Shield (Zymo Research #R1100-50), vortexed heavily, transferred to fresh PCR tubes, and shipped to Plasmidsaurus. RNAseq analysis was done by Plasmidsaurus and all RNAseq data figures were generated using their online interface.

### Flow cytometry

NL-CI^JRFL Env^-infected syncytia-producing cocultures (2x10^6^ total cells; containing a mixture of syncytia, infected mononucleated cells, and uninfected A3R5-G1D cells) were harvested, pelleted, and resuspended in PBS/0.1% bovine serum albumin (BSA). As a control, 5x10^5^ A301-G2D and A3R5.7-G1D cells were harvested and combined at a 1:1 ratio. Cells were pelleted and resuspended in PBS/0.1% BSA and stained with LIVE/DEAD Fixable Blue (Thermo Scientific #L34962) for 30 min on ice. Cells were then pelleted and resuspended in PBS/0.1% BSA containing an antibody cocktail (Table 1) and incubated on ice for 30 min in the dark. Cells were pelleted, resuspended, and fixed using PBS/4% paraformaldehyde (PFA; Electron Microscopy Sciences #50-980-494) for 10 min at room temperature in the dark, before washing and storing in PBS at 4°C for flow cytometry analysis.

**Table 1.** Flow cytometry antibody cocktail.

| Antibody/Reagent | Catalog # | Amount used |
| --- | --- | --- |
| Super Bright Complete Staining Buffer | eBioscience #SB440142 | 1:20 dilution |
| CD81-Brilliant Violet 421 | BD Biosciences #740079 | 5 µg/mL |
| HLA-B-CF405M | Biotum #75950-826 | 5 µg/mL |
| CD44-BUV496 | BD Biosciences #750519 | 2.5 µg/mL |
| CD155-PE | Biolegend # 337610 | 625 ng/mL |
| ULBP2/5/6 Brilliant Violet 605 | BD Biosciences #748131 | 10 µg/ml |
| HLA-F (custom conjugated with PE-CY5.5) | Biolegend #373202 | 1:1280 dilution |
| HLA-A-R718 | BD Biosciences #568031 | 6.25 µg/mL |
| CD4-BUV737 | BD Biosciences #612748 | 312.5 ng/mL |
| CD112-PE-CY7 | Biolegend #337414 | 5 µg/mL |
| ULBP3-Alexa Fluor 750 | R&D Systems #FAB1517S | 10 µg/mL |
| CD352 (custom conjugated with Alexa Fluor 790) | R&D Systems #MAB19081 | 1:160 dilution |
| CD3-BUV805 | Invitrogen #368-0037-42 | 250 ng/mL |

Samples were analyzed on a 4-laser Cytek Aurora (UV-V-B-R) flow cytometer, and data analysis was performed using FlowJo v10 (BD Biosciences) and the FlowJo UMAP plugin (version 4.1.1). UMAP dimensionality reduction was performed on cells that were first gated as Live (LIVE/DEAD Fixable Blue-negative), and the LIVE/DEAD Fixable Blue, mCherry, and EGFP parameters were excluded from the clustering process completely, leaving only the surface marker antibodies, none of which directly report whether a cell is uninfected, infected, or syncytial. After UMAP, ground-truth gates were established for uninfected cells (mCherry-negative, EGFP-negative), infected mononucleated (mCherry-positive, EGFP-negative), and syncytia (mCherry-positive, EGFP-positive). These were then were used to subset the data and investigate the clustering pattern for each cell category when plotting UMAP components, as well as to generate single-parameter histogram plots for each antibody label.

### 2D Killing experiments

KHYG-1 cells (hereafter referred to as “Effector cells”) were harvested, pelleted and resuspended in CellTrace Far Red (1:15,000 dilution) (Invitrogen #C34564) in PBS/100 U/mL IL2 at 1x10^6^ cells/mL. After 30 min at 37°C/5% CO_2_, unreacted dye was quenched by adding an equal volume of complete RPMI medium. Dyed KHYG-1 cells were pelleted and resuspended at a density of 1.8x10^6^ cells/mL in complete RPMI+IL-2. Syncytia-producing cocultures (hereafter referred to as “Target cells”) were pelleted and resuspended at a density of 1x10^6^ cells/mL in complete RPMI+IL-2.

1.5x10^5^ Target cells and 4.5x10^5^ Effector cells (1T:3E ratio) were combined and plated in a 48-well plate at 1.5x10^6^ cells/mL in triplicate. In addition, 1.5x10^5^ Target cells and 2.5x10^5^ Effector cells were cultured separate from each other in triplicate. Cells were incubated at 37°C/5% CO_2_ for 6 h.

After 6 h, cocultured cells were harvested and non-cocultured controls were harvested and combined. Cells were pelleted and resuspended in PBS/0.1% BSA and stained with LIVE/DEAD Fixable Blue (Thermo Scientific #L34962) for 30 min on ice. Cells were then pelleted and fixed using PBS/4% PFA for 10 min at room temperature in the dark, before washing and storing in PBS at 4°C for flow cytometry analysis.

### 3D Killing assay

KHYG-1 cells (“Effector cells”) were harvested, pelleted and resuspended in CellTrace Far Red (1:15,000 dilution) (Invitrogen #C34564) in PBS/100 U/mL IL2 at 5x10^7^ cells/mL. Syncytia-producing cocultures (“Target cells”) were pelleted and resuspended in complete RPMI+IL-2 at 5x10^7^ cells/mL.

PureCol (bovine Collagen I, ∼3 mg/mL solution) (Advanced Biomatrix #5005) was combined with 10X MEM (Gibco #11-430-030), sodium bicarbonate (7.5%; Gibco #25080-094), and 100 U/mL IL-2 on ice. For each 60 μL gel, 7.5x10^4^ Target cells were combined with 2.25x10^5^ Effector cells (1T:3E ratio) and then added to the gel mixture, yielding a final collagen concentration of 1.6 mg/mL, and ∼2.5% of the gel volume occupied by cells. Additionally, 7.5x10^4^ Target cells and 2.25x10^5^ Effector cells were cultured separately from the other cell type in collagen gels of proportionally smaller volume to maintain the same gel density. Gels were deposited in the center of the wells in a 24-well plate and allowed to set for 1 h at 37°C/5% CO_2_. Gels were then immersed in complete RPMI+IL-2. After 5 additional hours of incubation, supernatants were aspirated, and gels were dissolved using 1 mg/mL collagenase (Sigma-Aldrich #C0130-100MG) in RPMI-1640 and manually dissociated by pipetting. Cells were then washed, stained with LIVE/DEAD Fixable Blue on ice for 30 min, fixed in PBS/4% PFA for 10 min, and stored in PBS for flow cytometry.

### Flow cytometry for 2D and 3D killing assays

Samples were analyzed either on a 4-laser Cytek Aurora (UV-V-B-R) flow cytometer, or a 5-laser BD FACSDiscover S8 flow sorter (without sorting). Data analysis was performed using FlowJo v10 (BD Biosciences). Data were first gated for live cells (LIVE/DEAD Blue-negative), then Effector cells (CellTrace Far Red-positive, mCherry-negative) or Target cells (CellTrace Far Red-negative, mCherry-positive). Target cells were subsequently gates for mononucleated cells (EGFP-negative) or syncytia (EGFP-positive). To quantify killing, target cell counts for each cell subset (uninfected cells, infected mononucleated cells, and syncytia) were first normalized to effector counts within the same sample, then averaged across 3-4 technical replicates. Those averaged normalized counts within cocultured samples were divided by the averaged normalized counts in non-coculture controls, yielding the normalized proportion of cells of each subset that were still alive after coculture. This value was then subtracted from 1 and expressed as a percentage to quantify the proportion of cells of each subset that were missing (i.e., killed cells). To compare the killing of syncytia to infected mononucleated cells across 3-4 biological replicates, a two-tailed paired *t* test was used in GraphPad Prism 11.

